# A cell-type-specific nitric oxide-cGMP pathway regulates proprioceptor morphology

**DOI:** 10.1101/2023.04.14.536921

**Authors:** Raphael Cohn, Natalie Kolba, Jaqueline A. Picache, Nancy Lin, Aisha Saleem, Wesley B. Grueber

## Abstract

Neurons show remarkable morphological diversity facilitated by transcriptional regulation during development and fitting with their connectivity and function. Second messenger pathways mediated by cyclic 3’,5’ adenosine or guanosine monophosphate (cAMP or cGMP) may serve to integrate extrinsic signals and regulate branching and growth of neurites. We investigated the regulation of cGMP signaling in neuronal morphogenesis in *Drosophila* somatosensory neurons and found cell type-specificity in expression of the enzyme soluble guanylate cyclase (sGC) in different sensory neuron classes. sGC is responsible for the production of cGMP in response to the diffusible free radical messenger nitric oxide (NO) such that different levels of sGC impart cell type-specific sensitivity to NO. Supporting a role for the NO-cGMP pathway in neuronal morphogenesis, we find that knockout of sGC or nitric oxide synthase (NOS) impacts the morphology of multidendritic proprioceptors. The homeodomain transcription factor Cut shows different levels of expression in different somatosensory neuron types and we find that Cut regulates the level of cGMP signaling via repression of sGC expression. Thus, transcription factor levels contribute to cell type diversification by regulating levels of a signaling pathway to mediate somatosensory neuron morphogenesis.

## Introduction

Neurons show striking morphological and functional diversity determined by both intrinsic pathways and extrinsic cues^1–3^. Whereas studies of intrinsic factors that determine dendrite morphology, including transcription factors, cell adhesion molecules, and actin regulators, are fairly advanced, we have much to learn about responses to factors released from nearby tissues that may have special roles in coordinating neuronal morphogenesis and changes in body development or physiology.

One essential signaling pathway that operates in a variety of tissues and in different contexts is the nitric oxide-cyclic GMP (NO-cGMP) pathway^4–9^. NO is a diffusible gaseous messenger that is produced in cells by NO synthase (NOS) and can diffuse past cell membranes to signal to neighboring cells^6,10,11^. One prominent target of NO is soluble guanylate cyclase (sGC), binding of which catalyzes the production of cGMP. As a second messenger cGMP can modulate the activity of cGMP-dependent protein kinases (PKGs), phosphodiesterases (PDEs), and ion channels^12,13^. Prior studies showed persistent expression of NO-cGMP signaling pathway components in moth sensory neurons, in particular in a class of subepidermal somatosensory neurons called dendritic arborization (da) neurons^14^. However, the neurodevelopmental or functional roles of this signaling cascade in the somatosensory system are unknown.

The somatosensory system of *Drosophila* larvae is particularly well-suited to study how signals from nearby tissues affect neuronal form and function. Prior anatomical analysis segregated the *Drosophila* da somatosensory neurons into four classes, class I-IV, in order of increasing dendritic branching complexity^15^ (**Figure 1A**). Each class of neuron is identifiable based on their characteristic morphology, and shows distinct molecular signatures, axon projection patterns, and functions^16–30^. da neuron dendrites extend along the epidermis in a largely two-dimensional and highly overlapping fashion. Substrate interactions that control growth and maintenance are mediated by a number of cell surface adhesion molecules^22,31–41^. How substrate-derived signals that are widely available to each class of da neuron may mediate class-specific neuronal morphology is still not well understood.

**Figure 1.**
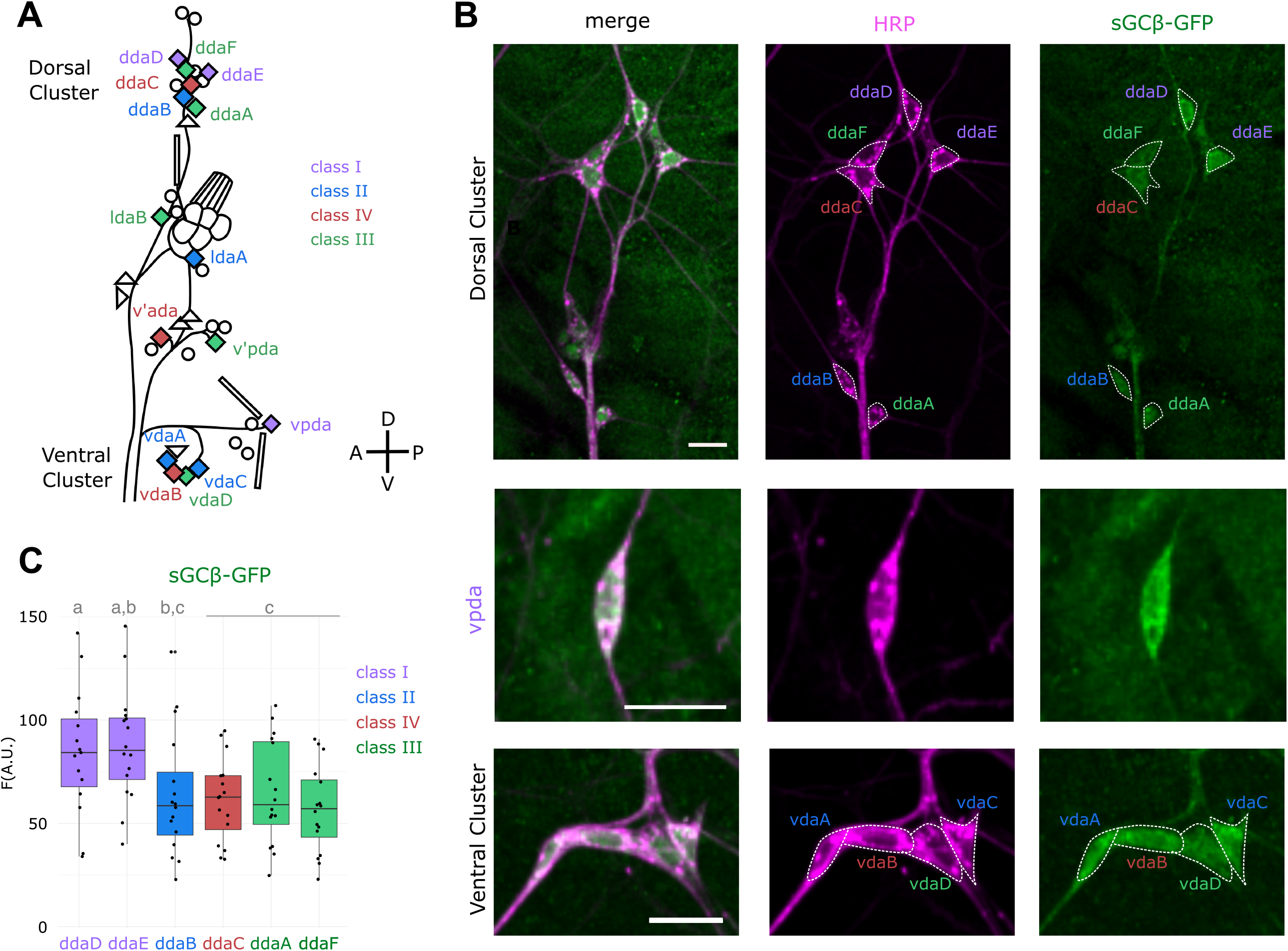
Sensory neuron classes express distinct levels of sGCβ. **A**, Diagram of peripheral sensory neurons present in each abdominal hemisegment of *Drosophila* larvae. Colored diamonds represent da neurons that are segregated into four distinct classes, as indicated by coloring. Circles represent external sensory neurons, triangles represent other md neurons and bars represent chordotonal organs. **B**, Representative images showing expression levels of GFP-tagged sGCβ (green) in dorsal and ventral cluster da neurons.**C**, Mean fluorescence intensities of GFP-tagged sGCβ in dorsal cluster da neurons. Here and throughout manuscript: HRP staining (magenta) highlights all dendrites and soma on the body wall; individual da soma are outlined with dashed lines; cell names are colored according to class as in **A**; scale bars in all representative images = 10um; lowercase letters above box plots indicate statistically significant differences between groups, with Bonferroni-corrected P-values < 0.05.

In this study, we show neuronal class-specific cGMP signaling in *Drosophila* somatosensory neurons. NO-induced cGMP production through sGC is high in the simplest, proprioceptive, class of da neurons, and loss of this pathway during development leads to defects in dendritic morphogenesis. We show that class-specificity of cGMP signaling is controlled by the homeodomain transcription factor Cut^17^. Altogether, we propose that a diffusible cue impacts cGMP signaling in a Cut-dependent class-specific manner to control dendritic arborization.

## RESULTS

### Evidence for functional NO-cGMP signaling in *Drosophila* somatosensory neurons

Prior results in Manduca sexta moths indicated a possible role for NO-cGMP signaling in somatosensory neurons. To examine roles for this pathway in *Drosophila* we first examined expression of an endogenously tagged soluble guanylate cyclase (sGCβ)-GFSTF protein^42^. We observed expression of tagged sGCβ in da neurons along the body wall, most prominently in the proprioceptive class I da neurons, with lower expression in mechanoreceptive and nociceptive sensory neurons (**Figure 1B-C**). sGC was present in sensory neuron cell bodies, dendrites and axons. There are three class I da neurons in each hemisegment and each was similarly positive for sGC expression (**Figure 1C**). These observations suggest that somatosensory neurons have the capacity to signal via cGMP, most prominently in proprioceptive neurons.

NO-sensitive sGC is a heterodimer of α and β sGC subunits, so expression of the β subunit alone is not sufficient for signaling. We took a pharmacological approach to determine whether fully constituted sGC was present in larval sensory neurons^43^. We exposed fileted third instar larvae to the NO donor, sodium nitroprusside (SNP) to stimulate sGC and the phosphodiesterase inhibitor 3-isobutyl-1-methylxanthine (IBMX) to prevent the breakdown of any cGMP that is produced. We detected significant cGMP production in class II and proprioceptive class I neurons and lower levels in other da sensory neurons (**Figure 2A-B**), suggesting that sGCα is expressed in da neurons, in addition to the observed expression of sGCβ. Each class I neuron showed comparable sensitivity to SNP+IBMX stimulation and we observed cGMP accumulation in cell bodies, dendrites, and axons. When we exposed homozygous *sGCα*^*253*^ mutants to SNP + IBMX we did not observe accumulation of cGMP (**Figure 2C**), indicating that SNP-sensitive cGMP production is mediated by sGC. Together these data suggest that da somatosensory neurons express components of an NO-sensitive sGC-cGMP signaling pathway in a cell type-specific manner.

**Figure 2.**
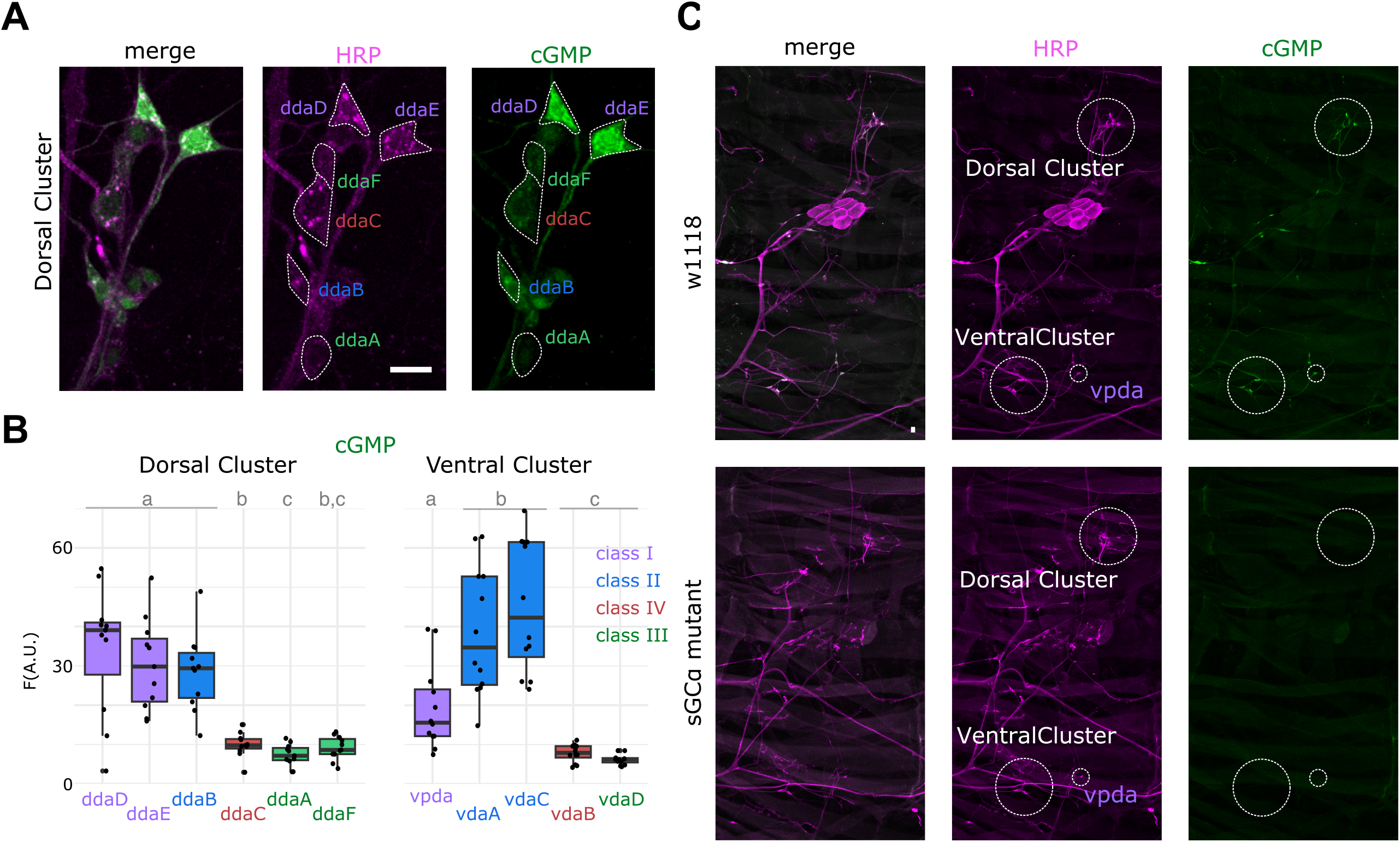
Sensory neuron classes generate distinct cGMP levels in response to NO stimulation. **A**, Representative images showing variation between dorsal cluster da cell classes of cGMP-antibody staining following exogenous NO stimulation with SNP-IBMX mix (see Methods). HRP highlights neuronal anatomy. **B**, Mean cGMP-staining fluorescence following NO stimulation measured in dorsal cluster (left) and ventral cluster (right). **C**, Representative images of NO-stimulated cGMP production throughout an entire abdominal hemisegment in control (*w*^*1118*^, top) and homozygous sGC mutant (*sGC*^*253*^, bottom) larvae. sGC mutation eliminates detectable cGMP response in all sensory neurons.

### Endogenous activation of cGMP signaling in sensory neurons

Activation of cGMP production in the above assays depends on an exogenous supply of NO and so does not reveal whether NO is an endogenous ligand for sGC in sensory neurons. To address whether endogenous NO is available to somatosensory neurons we dissected larvae and incubated them in IBMX alone to prevent breakdown of cGMP. We found that addition of IBMX alone post-dissection resulted in more modest, but detectable levels of cGMP accumulation in sensory neurons, with the highest levels in the class I neurons ddaD and ddaE (**Figure 3A-B**). To confirm cGMP production was due to NOS activity we performed the same experiment with addition of the NOS inhibitor L-nitroarginine methylester (L-NAME) and found a significant reduction of endogenous cGMP immunoreactivity (**Figure 3B**). By contrast, addition of D-NAME, the relatively inactive enantiomer of L-NAME, led to no such reduction of endogenous cGMP production (**Figure 3B**). Given the short half-life of NO we speculate that nearby expressed NOS provides a source of NO that signals to da neurons.

**Figure 3.**
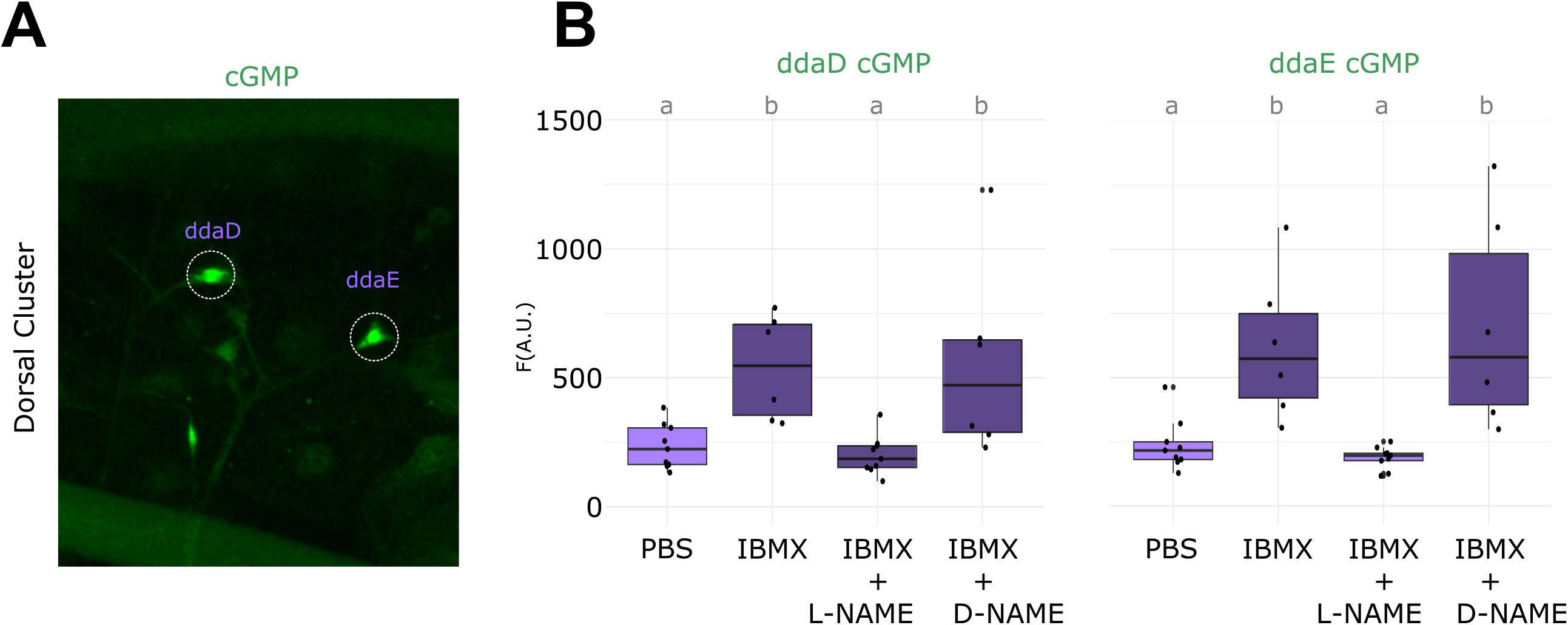
Evidence that endogenous NO release is sufficient to stimulate cGMP production in class I da neurons. **A**, Representative image of cGMP antibody staining in dorsal cluster following incubation of live dissected *w*^*1118*^ larvae with IBMX alone, without an exogenous NO donor. **B**, Mean intensity of cGMP immunofluorescence in class I neurons ddaD (left) and ddaE (right) after incubation in PBS alone (control), 1mM IBMX, IBMX+NOS-inhibitor L-NAME, and IBMX+inactive L-NAME enantiomer D-NAME.

### sGC controls proprioceptor dendrite morphology

Given our evidence for NO-cGMP signaling in class I sensory neurons in the peripheral nervous system, we explored potential effects on dendrite development. Class I neurons that show the strongest NO sensitivity are noteworthy for their relatively simple arborization that is polarized along the major axes of the larval body. Specifically, for each class I neuron a primary dendrite extends along the dorsoventral (DV) axis from which secondary branches extend along the anteroposterior (AP) axis (**Figure 4A-B**). Secondary dendrites are polarized by the extent of growth in anterior or posterior directions. We quantified several features of the third instar dendrite morphology of the class I neuron vpda in *sGCα*^*253*^ mutant larvae and found no obvious difference in the number of branch points or dendritic territory size (**Figure 4C**,**E**), but did find a difference in the polarity of secondary dendrites along the AP axis. Whereas in control larvae secondary dendrites showed more growth toward the posterior of the larva as measured by the ratio of total cable length of P-vs. A-directed secondary dendrites relative to the primary dendrite, in *sGCα*^*253*^ mutants dendrite growth was roughly equal in anterior and posterior directions (**Figure 4F**) due to an overall reduction in posterior branch length (p<0.0005). Animals heterozygous for *sGCα*^*253*^ and a deficiency containing the *sGCα* locus recapitulated this phenotype (**Figure 4F**). Similar secondary dendrite polarity defects were observed in animals unable to synthesize NO due to a mutation in NOS (**Figure 4D,G)**^44^.Taken together, these data suggest that dendrite morphology is regulated by NO-cGMP signaling.

**Figure 4.**
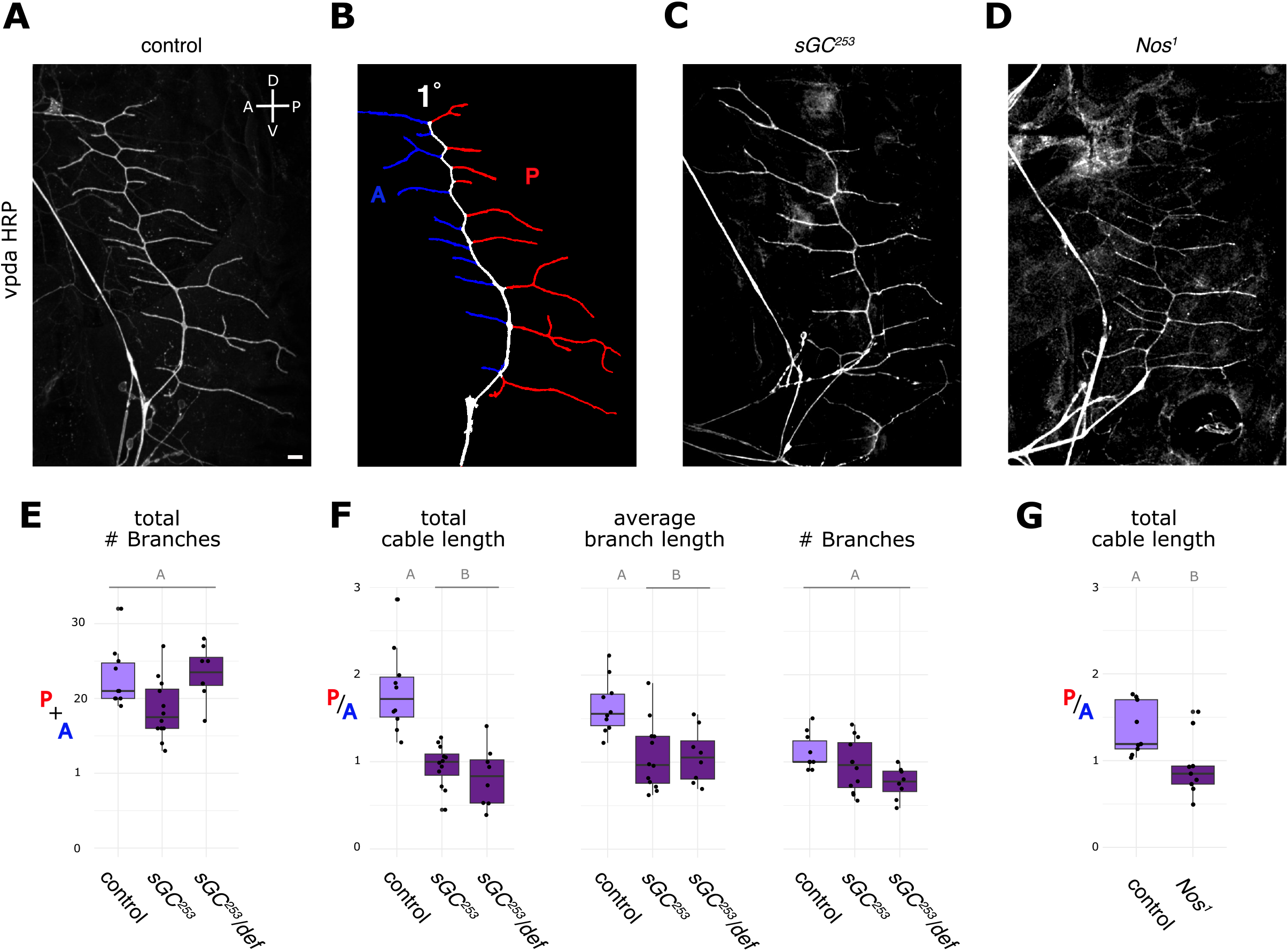
NO-cGMP signaling regulates vpda dendritic morphology. **A**, Representative image of HRP-stained vpda dendrites in *w*^*1118*^ control larvae. **B**, Diagram highlighting primary dendrite (white), total cable length measurement of anterior (A, blue) and posterior (P, red) secondary dendrites. **C**, Representative image of HRP-stained vpda dendrites in *sGC*^*253*^ mutants and **D**, in *Nos*^*1*^ mutants. **E**, Comparison of total number of secondary branches in *w*^*1118*^ (control), sGCɑ homozygous mutants (sGC^253^) and animals heterozygous for *sGC*^*253*^ and a deficiency chromosome (Df(3R)BSC501) that covers the *sGCα* gene locus. **F**, Ratio of posterior to anterior vpda secondary dendrites measured in total cable lengths, left, average branch length, center, and number of branches, right. **G**, Ratio of posterior to anterior vpda secondary dendrites measured in total cable lengths in control compared with *Nos*^*1*^ mutants.

### Cut transcription factor controls cell specific NO-sensitivity

We next asked how cell-type specific patterns of NO sensitivity in the somatosensory system are regulated. The homeodomain transcription factor Cut is expressed in different levels in different classes of da neurons^17^, which are negatively correlated with the degree of NO-sensitivity. Class I and II neurons, which express low or undetectable levels of Cut, showed strongest sensitivity to endogenous and exogenous NO. Other da neurons that show stronger Cut expression show lower sensitivity to exogenous NO (**Figure 5A-C**). Given the inverse relationship between Cut and NO sensitivity, we hypothesized that Cut-mediated repression may establish class-specific responses to NO. To test this scenario, we mis-expressed Cut in all da neurons using *109(2)80-Gal4* and stimulated with SNP+IBMX. Increasing Cut expression reduced NO sensitivity in class I da neurons (**Figure 5D-F**). Thus, Cut is sufficient to repress NO sensitivity. Furthermore, Cut repression of NO sensitivity likely occurs through regulation of sGCβ expression since driving *UAS-cut* with *109(2)80-Gal4* suppressed sGCβ-GFSTF expression (**Figure 5G**).

**Figure 5.**
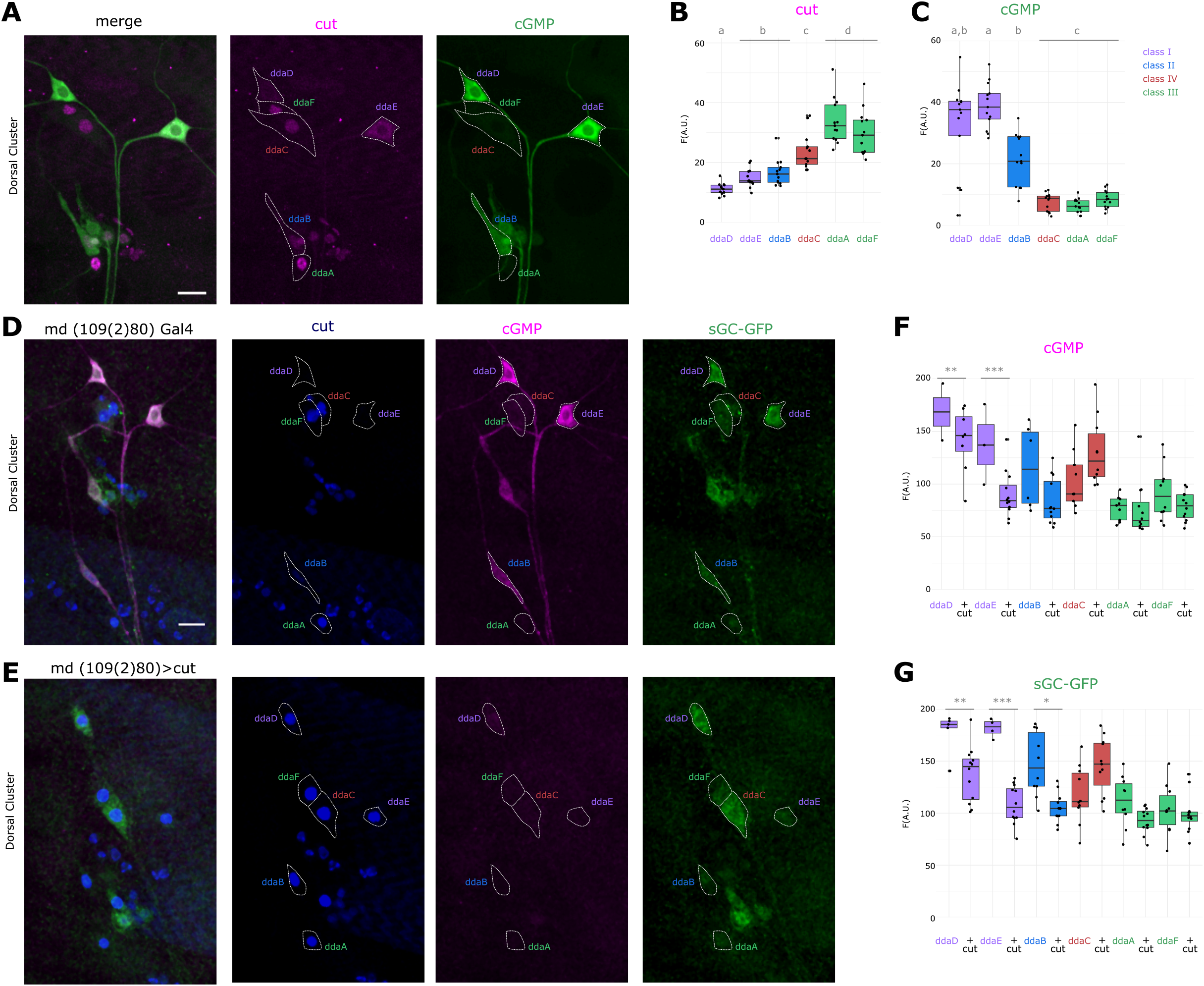
Cut transcription factor represses sGCβ expression and cGMP production in da neurons. **A**, Representative images of dorsal cluster neurons stained for Cut (magenta) and cGMP (green) following NO stimulation. **B**, Mean fluorescence intensity of nuclear cut (magenta) and **C**, somatic cGMP (green), in wildtype larvae following NO stimulation. **D**, Representative images showing mean immunofluorescence intensity of cut, cGMP and sGC-GFP in *109(2)80-Gal4* control animals and **E**, *109(2)80-Gal4* driving expression of *UAS-cut*^*5*^. **F**, Mean cGMP fluorescence intensity and **G**, Mean sGC-GFP fluorescence, for each dorsal cluster da neuron in control and cut-overexpression animals following NO stimulation. Asterisks indicate statistically significant comparisons after Bonferroni correction with p<0.05*, p<0.005**, and p<0.0005***.

We also tested whether Cut is necessary for the low sensitivity to NO shown by class III neurons using RNAi and CRISPR-mediated knockdown of Cut in class III neurons^45,46^. Both approaches led to loss of Cut immunostaining and loss of the characteristic class III dendritic branchlets, consistent with prior studies^17^ (**Figure 6A-B**). Reduction of Cut in class III neurons by either approach led to a significant increase in NO-mediated cGMP production (**Figure 6C,D,F**). Unexpectedly, relative cGMP responses in some class II and IV neurons also increased when CRISPR was used to knock down Cut using the class III driver 83B04-Gal4. Further investigation will be needed to determine whether these increases are due to off-target developmental expression or leakage of cas9 or cell non-autonomous effects of class III loss of Cut (**Figure 6E,G**). Taken together, these data suggest that Cut is both necessary and sufficient for suppression of NO-induced cGMP production in somatosensory neurons.

**Figure 6.**
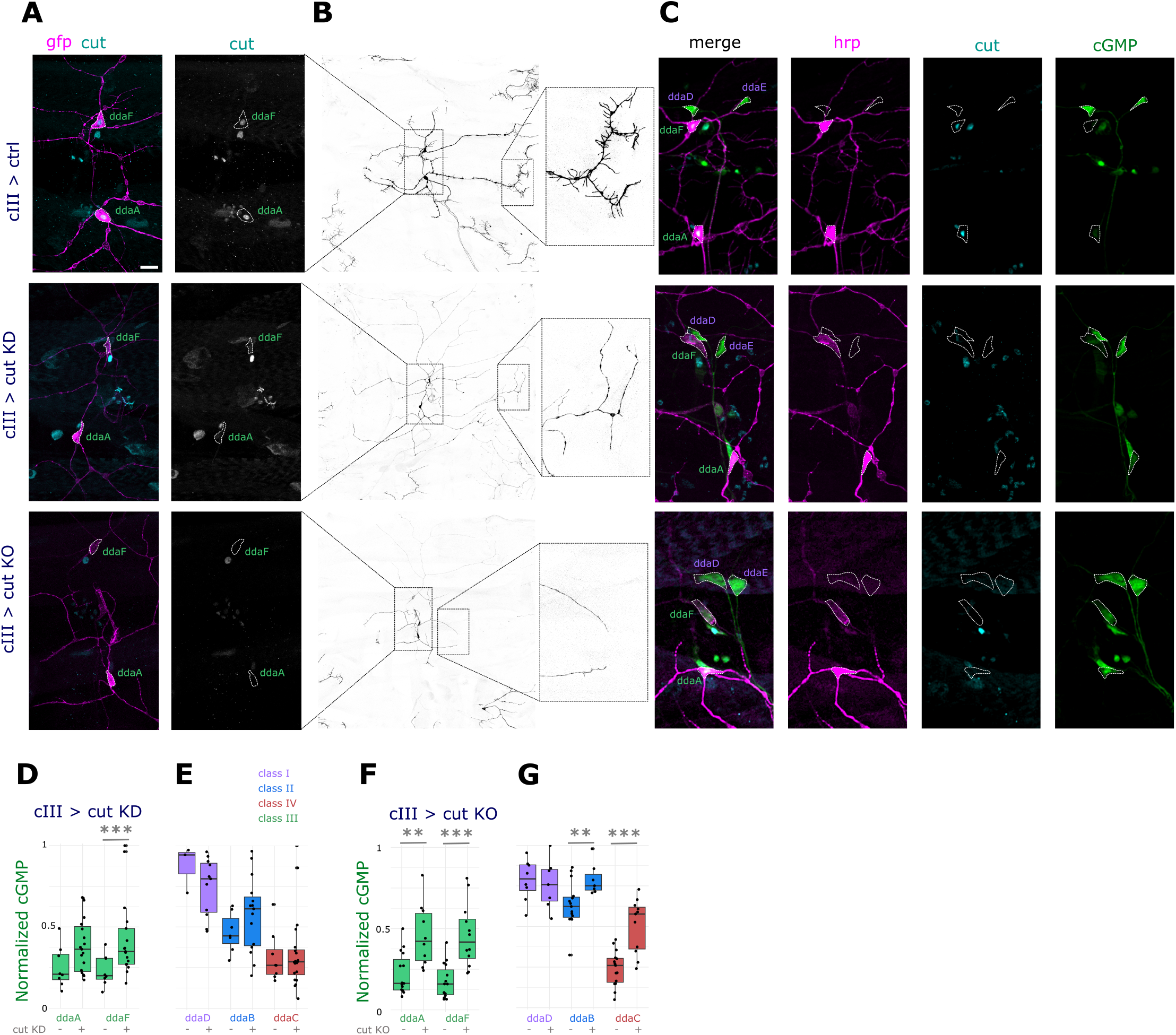
cut represses NO-induced cGMP productionin class III da neurons. **A**, Representative images showing dorsal cluster class III (cIII) neurons labeled with GFP (*83B04-Gal4, UAS-CD4tdGFP)* and nuclear cut immunofluorescence in control (ctrl; top), with *UAS-cutRNAi* (cut KD; middle) and with *UAS-cut-sgRNA* and *UAS-uMcas9* (cut KO; bottom). **B**, Zoomed out view and zoomed in insets showing loss of stereotypical class III dendrite morphology when cut is knocked down. **C**, Representative images showing cut and cGMP immunofluorescence after NO stimulation in the dorsal cluster class I and class III neurons in control (top) and cut-knockdown examples. **D**, Quantification of mean cGMP immunofluorescence in cIII neurons relative to ddaE, generally the class I neuron with the highest cGMP level. Comparison of control and cut RNAi. **E**, Same as **D** but for other da neuron classes not clearly targeted by *84B04-Gal4*. **F**,**G**, Same as **D**,**E** but comparing control and cut CRISPR.

## DISCUSSION

### NO-cGMP signaling in sensory neurons

In a pharmacological assay for NO-sensitive sGC activity we identified cell-type specific cGMP accumulation in *Drosophila* da sensory neurons. Consistent with this finding, we found similar cell-type specific expression of an sGC-linked reporter in somatosensory neuron soma. Maximal cGMP production required an exogenous source of NO but cGMP levels also increased endogenously if cGMP breakdown was blocked by bathing tissue in a phosphodiesterase inhibitor. This endogenous increase was also blocked by inhibitors of NOS, suggesting a nearby source of NO that signals to sensory neurons. Possible sources of NO include epidermal cells that provide a substrate for da neuron dendrites, body wall muscles, and glia that surround da neurons and proximal dendrite branches^31,32,40^.

In vertebrates, each of the different NOS isozymes (neuronal NOS, inducible NOS, and endothelial NOS) are expressed in the skin^4^. Roles are varied, including barrier formation, cell death, wound healing, inflammation, and responses to UV, however signaling to nearby somatosensory neurons has not, to our knowledge, been demonstrated. Given the many similarities between *Drosophila* and vertebrate somatosensory neurons in terms of morphology, gene expression, and function, roles for NO and cGMP signaling will be promising areas of study in other skin sensory systems.

### A role for cGMP in dendritic morphogenesis

Late third instar NO sensitivity of sensory neurons suggests ongoing developmental or functional roles for cGMP signaling. Class I da neurons, which are strongly sensitive to NO stimulation, extend dendrites in a polarized fashion along the anteroposterior axis. The vpda neuron, for example, extends secondary dendrites primarily in the posterior direction with a relatively minor anterior arborization^15^ In the absence of sGC dendrites arborized roughly equally in anterior and posterior directions. The class I vpda neuron elaborates by first extending a primary dendrite in a dorsal direction followed by extending roughly evenly spaced comb-like secondary dendrites in anterior and posterior directions^47^. This initial arborization is established by hatching, after which few new branches are formed but existing arbors expand to maintain scaling with the growing body wall^33^. We envision two possible scenarios for how the vpda dendrite phenotype could emerge. First, anterior and posterior-directed dendrites could grow to equal lengths in both directions, reflecting a defect in dendrite polarized growth. In this case, the positioning of the dendritic arbor would be shifted in an anterior direction. Second, the primary dendrite could take a more posterior position in the segment with no change in growth and positioning of secondary dendrites. In this case, the arbor would show an apparent shift in secondary dendrite polarity but the main defect would in fact be in primary dendrite location. An important future goal will be to determine the cues and timing that mediate polarized growth and how these are affected by cGMP signaling.

In other systems cGMP and cAMP mediate neuronal responses to developmental guidance cues such as Netrin and Semaphorin. The relative levels of cGMP and cAMP can determine attractive or repulsive responses to Netrin-1^48^. sGC localization to the base of the apical dendrite of pyramidal cells mediates attraction to Semaphorin 3A signals^49^, so it is possible that cGMP mediates dendritic arbor polarity or primary dendrite targeting by transducing or modulating the response to an extrinsic cue. Our data indicate that the morphology of vpda is under tight regulation, suggesting functional relevance to its dendritic morphology. Prior studies showed that posterior-directed vpda dendrites undergo the greatest deformation with each forward peristaltic wave during crawling^50,51^. Thus, the extent of anterior and posterior arborization of class I neurons may influence dendrite deformation and sensory response properties during locomotion.

### Cut regulation of cGMP signaling levels

Homeodomain transcription factors in the Cut family control neuronal morphogenesis in both vertebrate and invertebrate systems^17,21,52–55^. In *Drosophila* da sensory neurons, Cut is expressed at different levels in different morphological classes, with the simplest class I neurons lacking Cut and other classes expressing successively higher levels from cII < cIV < cIII. Increasing or decreasing Cut levels in different classes causes morphological switches toward the morphology of cells with higher or lower Cut levels, respectively. Thus, increases in Cut lead to more complex cIII-type morphologies, and reduction of Cut causes dendrite simplification. One unanswered question is how Cut exerts effects on neuronal form or function through its transcriptional targets. A number of possible downstream targets of Cut have been identified^21,54,56^. We observed an inverse correlation between Cut levels and the degree of NO-sensitivity of da neuron classes, fitting with a role for Cut proteins as transcriptional repressors^57–60^. Misexpression and loss of function manipulations supported a causal relationship between Cut levels and sGC activity. These results reveal that Cut levels in sensory neurons are linked to repression of gene expression levels and implicate cGMP signaling in neuronal morphogenesis. Additional studies to address the contribution of cGMP to Cut-dependent dendritic morphogenesis, and pathways downstream of cGMP, are important future goals.

## ACKNOWLEDGMENTS

We thank Drs. Jan De Vente and James Truman for generously sharing anti-cGMP and Dr. Yuh-Nung Jan for supporting early work on this project. We thank Luke Nunnelly for early work on this project. Research reported in this publication was supported by the National Institute of Neurological Disorders and Stroke of the National Institutes of Health under Award Number R01NS061908 (W.B.G.). R.M.C. was supported by a Simons Society of Fellows Junior Fellowship. The content is solely the responsibility of the authors and does not necessarily represent the official views of the National Institutes of Health.

## METHODS

### Experimental animals

*Drosophila* stocks were raised on standard media at 18 or 25°C. The following stocks were used: (1) *109(2)80-Gal4*^61^ (BDSC_8769), (2) *UAS-cut(5)*^62^, (3) *2-21-Gal4*^17^, (4) UAS-Dicer2 (BDSC_24644), (5) *sGC*^*253* 9^; (6) UAS-cutRNAi (generated in the laboratory of Dr. Yuh-Nung Jan); (7) *w*^*1118*^, sGCβ-GFP(y1w*;Mi{PT-GFSTF.2}Gycβ100BMI08892-GFSTF.2/TM6C, Sb^1^ Tb+)^63^ (BL_60565); (7) *UAS-mCD8-GFP*^64^ (BDSC_5137), (8) *83B04-Gal4*^65^ (BDSC_41309) (Galindo, *in press*), (9) Df(3R)BSC501 (sGCα deficiency; BDSC_25005) (10) *Nos*^*1*^; (11) UAS-uScas9 (VDRC_ID_340000), (12) Cut-sgRNA (VDRC ID 341662)^45,46^

### Nitric Oxide Stimulation

NO stimulation was performed essentially as described^14^. Live dissected third instar larvae were pinned and exposed to 5 mM Sodium nitroprusside (SNP, Sigma) and 1 mM 3-Isobutyl-1-methylxanthine (IBMX, Sigma) in PBS for 15 minutes, fixed in 4% PFA in PBS for 15 minutes, then washed with PBS-TX for 2 5-minute intervals. cGMP production was detected using sheep anti-cGMP (1:10,000) (Antibody courtesy of Drs. Jan De Vente and James Truman)^66^.

### Immunohistochemistry

Immunostaining was performed essentially as described^15^. Other primary antibodies used were goat anti-HRP (1:100), mouse anti-Cut (1:20), and chicken anti-GFP (1:1000). All secondary antibodies were from Donkey and diluted at 1:200 (Jackson Immunoresearch).

### Confocal Imaging

Imaging was performed on a Zeiss 700 or Zeiss 710 confocal microscope. Identical image collection settings (laser intensity, detector gain, scan settings) were used for all images within each dataset.

### Analysis

Image analysis was performed in FIJI using custom ImageJ macros. For each cell analyzed, the user manually selected the soma and nucleus in the Z-slice in which the cell/nucleus was brightest. The script then measured the mean intensity of each ROI in each fluorescence channel and output to CSV file. Exported data was analyzed using Excel and custom scripts in Rstudio, written with assistance from ChatGPT. Statistical analysis and graphs were generated in R. Each statistical comparison was tested by ANOVA followed by t-tests with Bonferroni correction for multiple comparisons. Tracing of vpda dendrites was performed using the Simple Neurite Tracer FIJI plugin. Representative images shown were Z-projected and display min/max was adjusted to maximize dynamic range while avoiding saturation as much as possible. Identical brightness settings were used for all channels/images being directly compared within each figure.

## Notes

### Competing Interest Statement

The authors have declared no competing interest.

## References

1. Dong, X., Shen, K. & Bülow, H. E. Intrinsic and extrinsic mechanisms of dendritic morphogenesis. Annu. Rev. Physiol. 77, 271–300 (2015).

2. Corty, M. M., Matthews, B. J. & Grueber, W. B. Molecules and mechanisms of dendrite development in Drosophila. Development 136, 1049–1061 (2009).

3. Lefebvre, J. L. Molecular mechanisms that mediate dendrite morphogenesis. Curr. Top. Dev. Biol. 142, 233–282 (2021).

4. Cals-Grierson, M.-M. & Ormerod, A. D. Nitric oxide function in the skin. Nitric Oxide 10, 179–193 (2004).

5. Wildemann, B. & Bicker, G. Nitric oxide and cyclic GMP induce vesicle release at Drosophila neuromuscular junction. J. Neurobiol. 39, 337–346 (1999).

6. Michel, T. & Vanhoutte, P. M. Cellular signaling and NO production. Pflugers Arch. 459, 807–816 (2010).

7. Steinert, J. R., Chernova, T. & Forsythe, I. D. Nitric oxide signaling in brain function, dysfunction, and dementia. Neuroscientist 16, 435–452 (2010).

8. Kalyanaraman, H., Schall, N. & Pilz, R. B. Nitric oxide and cyclic GMP functions in bone. Nitric Oxide 76, 62–70 (2018).

9. Gibbs, S. M., Becker, A., Hardy, R. W. & Truman, J. W. Soluble guanylate cyclase is required during development for visual system function in Drosophila. J. Neurosci. 21, 7705–7714 (2001).

10. Cooke, R. M., Mistry, R., Challiss, R. A. J. & Straub, V. A. Nitric oxide synthesis and cGMP production is important for neurite growth and synapse remodeling after axotomy. J. Neurosci. 33, 5626–5637 (2013).

11. Regulski, M. & Tully, T. Molecular and biochemical characterization of dNOS: a Drosophila Ca2+/calmodulin-dependent nitric oxide synthase. Proc. Natl. Acad. Sci. U. S. A. 92, 9072–9076 (1995).

12. Francis, S. H., Busch, J. L., Corbin, J. D. & Sibley, D. cGMP-dependent protein kinases and cGMP phosphodiesterases in nitric oxide and cGMP action. Pharmacol. Rev. 62, 525–563 (2010).

13. MacPherson, M. R., Lohmann, S. M. & Davies, S.-A. Analysis of Drosophila cGMP-dependent protein kinases and assessment of their in vivo roles by targeted expression in a renal transporting epithelium. J. Biol. Chem. 279, 40026–40034 (2004).

14. Grueber, W. B. & Truman, J. W. Development and organization of a nitric-oxide-sensitive peripheral neural plexus in larvae of the moth, Manduca sexta. J. Comp. Neurol. 404, 127–141 (1999).

15. Grueber, W. B., Jan, L. Y. & Jan, Y. N. Tiling of the Drosophila epidermis by multidendritic sensory neurons. Development 129, 2867–2878 (2002).

16. Grueber, W. B. et al. Projections of Drosophila multidendritic neurons in the central nervous system: links with peripheral dendrite morphology. Development 134, 55–64 (2007).

17. Grueber, W. B., Jan, L. Y. & Jan, Y. N. Different levels of the homeodomain protein cut regulate distinct dendrite branching patterns of Drosophila multidendritic neurons. Cell 112, 805–818 (2003).

18. Emoto, K. et al. Control of dendritic branching and tiling by the Tricornered-kinase/Furry signaling pathway in Drosophila sensory neurons. Cell 119, 245–256 (2004).

19. Emoto, K., Parrish, J. Z., Jan, L. Y. & Jan, Y.-N. The tumour suppressor Hippo acts with the NDR kinases in dendritic tiling and maintenance. Nature 443, 210–213 (2006).

20. Long, H., Ou, Y., Rao, Y. & van Meyel, D. J. Dendrite branching and self-avoidance are controlled by Turtle, a conserved IgSF protein in Drosophila. Development Vol. 136 3475–3484 Preprint at https://doi.org/10.1242/dev.040220 (2009).

21. Sulkowski, M. J., Iyer, S. C., Kurosawa, M. S., Iyer, E. P. R. & Cox, D. N. Turtle functions downstream of Cut in differentially regulating class specific dendrite morphogenesis in Drosophila. PLoS One 6, e22611 (2011).

22. Parrish, J. Z., Xu, P., Kim, C. C., Jan, L. Y. & Jan, Y. N. The microRNA bantam functions in epithelial cells to regulate scaling growth of dendrite arbors in drosophila sensory neurons. Neuron 63, 788–802 (2009).

23. Parrish, J. Z., Kim, M. D., Jan, L. Y. & Jan, Y. N. Genome-wide analyses identify transcription factors required for proper morphogenesis of Drosophila sensory neuron dendrites. Genes Dev. 20, 820–835 (2006).

24. Lee, J., Peng, Y., Lin, W.-Y. & Parrish, J. Z. Coordinate control of terminal dendrite patterning and dynamics by the membrane protein Raw. Development 142, 162–173 (2015).

25. Lin, W.-Y. et al. The SLC36 transporter Pathetic is required for extreme dendrite growth in Drosophila sensory neurons. Genes Dev. 29, 1120–1135 (2015).

26. Hu, C. et al. Conserved Tao Kinase Activity Regulates Dendritic Arborization, Cytoskeletal Dynamics, and Sensory Function in Drosophila. J. Neurosci. 40, 1819–1833 (2020).

27. Petersen, M., Tenedini, F., Hoyer, N., Kutschera, F. & Soba, P. Assaying Thermo-nociceptive Behavior in Drosophila Larvae. BIO-PROTOCOL Vol. 8 Preprint at https://doi.org/10.21769/bioprotoc.2737 (2018).

28. Tracey, W. D., Jr, Wilson, R. I., Laurent, G. & Benzer, S. painless, a Drosophila gene essential for nociception. Cell 113, 261–273 (2003).

29. Honjo, K., Mauthner, S. E., Wang, Y., Skene, J. H. P. & Tracey, W. D., Jr. Nociceptor-Enriched Genes Required for Normal Thermal Nociception. Cell Rep. 16, 295–303 (2016).

30. Stewart, A., Tsubouchi, A., Rolls, M. M., Tracey, W. D. & Sherwood, N. T. Katanin p60-like1 promotes microtubule growth and terminal dendrite stability in the larval class IV sensory neurons of Drosophila. J. Neurosci. 32, 11631–11642 (2012).

31. Han, C. et al. Integrins regulate repulsion-mediated dendritic patterning of drosophila sensory neurons by restricting dendrites in a 2D space. Neuron 73, 64–78 (2012).

32. Kim, M. E., Shrestha, B. R., Blazeski, R., Mason, C. A. & Grueber, W. B. Integrins establish dendrite-substrate relationships that promote dendritic self-avoidance and patterning in drosophila sensory neurons. Neuron 73, 79–91 (2012).

33. Parrish, J. Z., Emoto, K., Kim, M. D. & Jan, Y. N. Mechanisms that regulate establishment, maintenance, and remodeling of dendritic fields. Annu. Rev. Neurosci. 30, 399–423 (2007).

34. Jiang, N., Soba, P., Parker, E., Kim, C. C. & Parrish, J. Z. The microRNA bantam regulates a developmental transition in epithelial cells that restricts sensory dendrite growth. Development 141, 2657–2668 (2014).

35. Jiang, N. et al. A conserved morphogenetic mechanism for epidermal ensheathment of nociceptive sensory neurites. Elife 8, (2019).

36. Meltzer, S. et al. Epidermis-Derived Semaphorin Promotes Dendrite Self-Avoidance by Regulating Dendrite-Substrate Adhesion in Drosophila Sensory Neurons. Neuron 89, 741–755 (2016).

37. Tenenbaum, C. M., Misra, M., Alizzi, R. A. & Gavis, E. R. Enclosure of Dendrites by Epidermal Cells Restricts Branching and Permits Coordinated Development of Spatially Overlapping Sensory Neurons. Cell Rep. 20, 3043–3056 (2017).

38. Yang, W.-K. & Chien, C.-T. Beyond being innervated: the epidermis actively shapes sensory dendritic patterning. Open Biol. 9, 180257 (2019).

39. Yang, W.-K. et al. Epidermis-Derived L1CAM Homolog Neuroglian Mediates Dendrite Enclosure and Blocks Heteroneuronal Dendrite Bundling. Curr. Biol. 29, 1445–1459.e3 (2019).

40. Han, C., Jan, L. Y. & Jan, Y.-N. Enhancer-driven membrane markers for analysis of nonautonomous mechanisms reveal neuron–glia interactions in Drosophila. Proceedings of the National Academy of Sciences Vol. 108 9673–9678 Preprint at https://doi.org/10.1073/pnas.1106386108 (2011).

41. Hoyer, N. et al. Ret and Substrate-Derived TGF-β Maverick Regulate Space-Filling Dendrite Growth in Drosophila Sensory Neurons. Cell Rep. 24, 2261–2272.e5 (2018).

42. Nagarkar-Jaiswal, S. et al. A genetic toolkit for tagging intronic MiMIC containing genes. Elife 4, (2015).

43. Truman, J. W., De Vente, J. & Ball, E. E. Nitric oxide-sensitive guanylate cyclase activity is associated with the maturational phase of neuronal development in insects. Development 122, 3949–3958 (1996).

44. Lacin, H. et al. Genome-wide identification of Drosophila Hb9 targets reveals a pivotal role in directing the transcriptome within eight neuronal lineages, including activation of nitric oxide synthase and Fd59a/Fox-D. Dev. Biol. 388, 117–133 (2014).

45. Meltzer, H. et al. Tissue-specific (ts)CRISPR as an efficient strategy for in vivo screening in Drosophila. Nat. Commun. 10, 2113 (2019).

46. Port, F. et al. A large-scale resource for tissue-specific CRISPR mutagenesis in Drosophila. Elife 9, (2020).

47. Sugimura, K. et al. Distinct developmental modes and lesion-induced reactions of dendrites of two classes of Drosophila sensory neurons. J. Neurosci. 23, 3752–3760 (2003).

48. Nishiyama, M. et al. Cyclic AMP/GMP-dependent modulation of Ca2+ channels sets the polarity of nerve growth-cone turning. Nature 423, 990–995 (2003).

49. Polleux, F., Morrow, T. & Ghosh, A. Semaphorin 3A is a chemoattractant for cortical apical dendrites. Nature 404, 567–573 (2000).

50. Vaadia, R. D. et al. Characterization of Proprioceptive System Dynamics in Behaving Drosophila Larvae Using High-Speed Volumetric Microscopy. Curr. Biol. 29, 935–944.e4 (2019).

51. He, L. et al. Direction Selectivity in Drosophila Proprioceptors Requires the Mechanosensory Channel Tmc. Curr. Biol. 29, 945–956.e3 (2019).

52. Bodmer, R. et al. Transformation of sensory organs by mutations of the cut locus of D. melanogaster. Cell 51, 293–307 (1987).

53. Cubelos, B. et al. Cux1 and Cux2 regulate dendritic branching, spine morphology, and synapses of the upper layer neurons of the cortex. Neuron 66, 523–535 (2010).

54. Iyer, S. C. et al. Cut, via CrebA, transcriptionally regulates the COPII secretory pathway to direct dendrite development in Drosophila. J. Cell Sci. 126, 4732–4745 (2013).

55. Iyer, E. P. R. et al. Functional genomic analyses of two morphologically distinct classes of Drosophila sensory neurons: post-mitotic roles of transcription factors in dendritic patterning. PLoS One 8, e72434 (2013).

56. Bhattacharya, S., Iyer, E. P. R., Iyer, S. C. & Cox, D. N. Cell-type specific transcriptomic profiling to dissect mechanisms of differential dendritogenesis. Genom Data 2, 378–381 (2014).

57. Dufort, D. & Nepveu, A. The human cut homeodomain protein represses transcription from the c-myc promoter. Mol. Cell. Biol. 14, 4251–4257 (1994).

58. Mailly, F. et al. The human cut homeodomain protein can repress gene expression by two distinct mechanisms: active repression and competition for binding site occupancy. Mol. Cell. Biol. 16, 5346–5357 (1996).

59. Stratigopoulos, G., LeDuc, C. A., Cremona, M. L., Chung, W. K. & Leibel, R. L. Cut-like homeobox 1 (CUX1) regulates expression of the fat mass and obesity-associated and retinitis pigmentosa GTPase regulator-interacting protein-1-like (RPGRIP1L) genes and coordinates leptin receptor signaling. J. Biol. Chem. 286, 2155–2170 (2011).

60. Nepveu, A. Role of the multifunctional CDP/Cut/Cux homeodomain transcription factor in regulating differentiation, cell growth and development. Gene 270, 1–15 (2001).

61. Gao, F. B., Brenman, J. E., Jan, L. Y. & Jan, Y. N. Genes regulating dendritic outgrowth, branching, and routing in Drosophila. Genes Dev. 13, 2549–2561 (1999).

62. Blochlinger, K., Jan, L. Y. & Jan, Y. N. Transformation of sensory organ identity by ectopic expression of Cut in Drosophila. Genes Dev. 5, 1124–1135 (1991).

63. Nagarkar-Jaiswal, S. et al. A library of MiMICs allows tagging of genes and reversible, spatial and temporal knockdown of proteins in Drosophila. Elife 4, (2015).

64. Lee, T. & Luo, L. Mosaic analysis with a repressible cell marker for studies of gene function in neuronal morphogenesis. Neuron 22, 451–461 (1999).

65. Jenett, A. et al. A GAL4-driver line resource for Drosophila neurobiology. Cell Rep. 2, 991–1001 (2012).

66. de Vente, J., Steinbusch, H. W. & Schipper, J. A new approach to immunocytochemistry of 3’,5’-cyclic guanosine monophosphate: preparation, specificity, and initial application of a new antiserum against formaldehyde-fixed 3’,5’-cyclic guanosine monophosphate. Neuroscience 22, 361–373 (1987).

